## Supplementary Figures and Tables for "TRIM7 restricts Coxsackievirus and norovirus infection by detecting the C-terminal glutamine generated by 3C protease processing"

**Figure S1**. ITC traces for GYG protein and main peptides

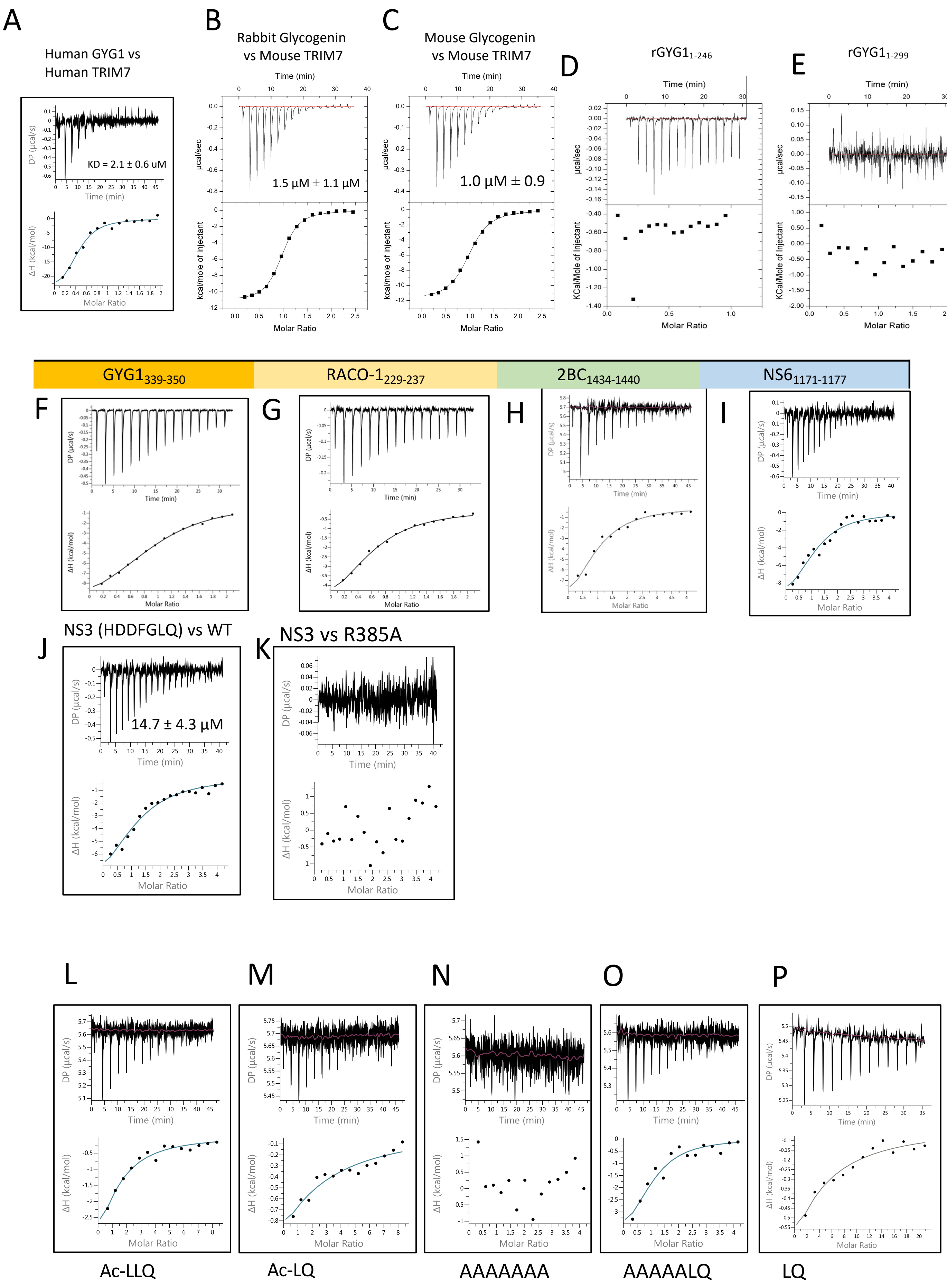

### **Figure S2.** Crystal structures of TRIM7-peptide complexes

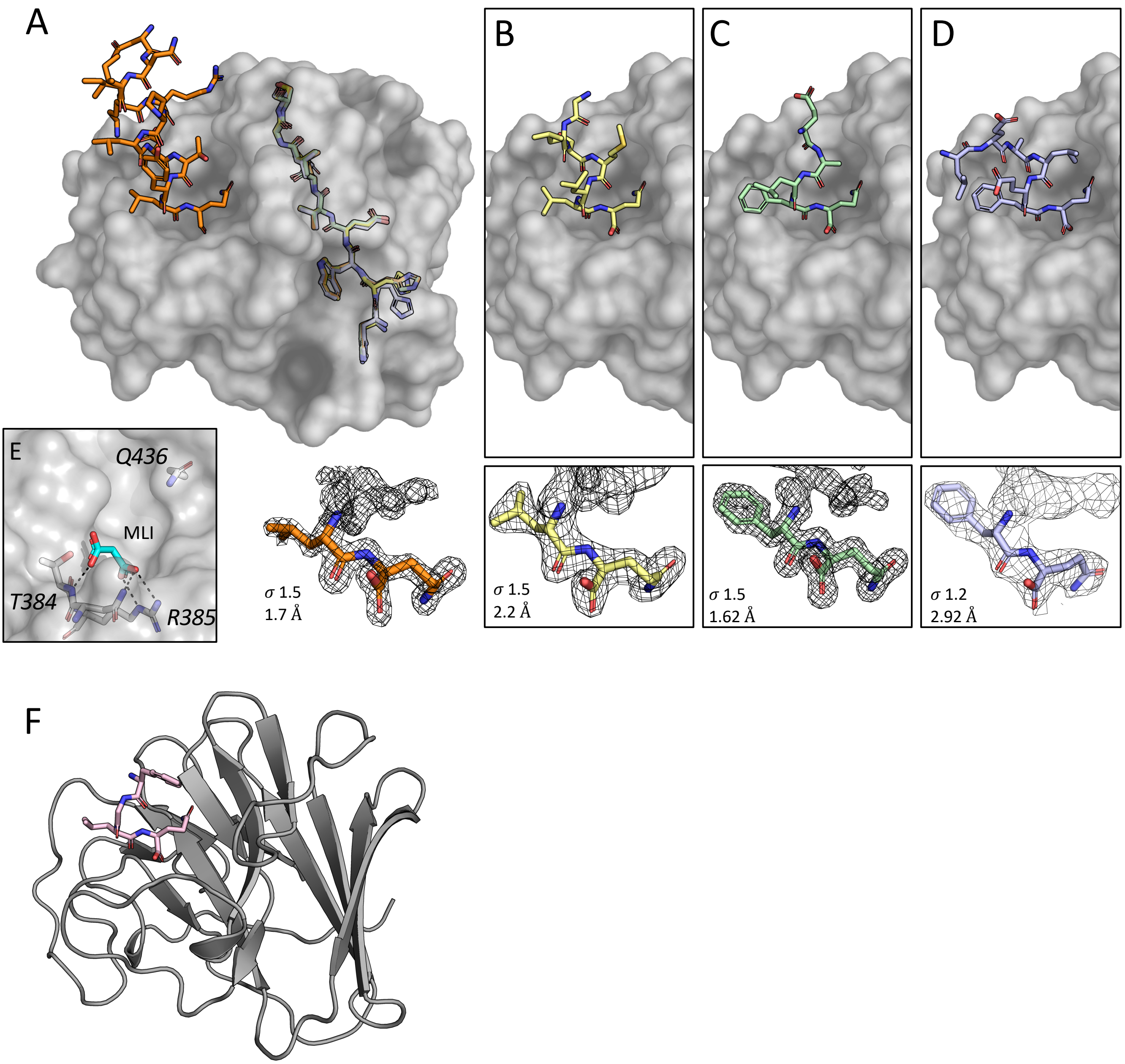

### **Figure S3** ITC traces for mutant TRIM7 proteins and peptides

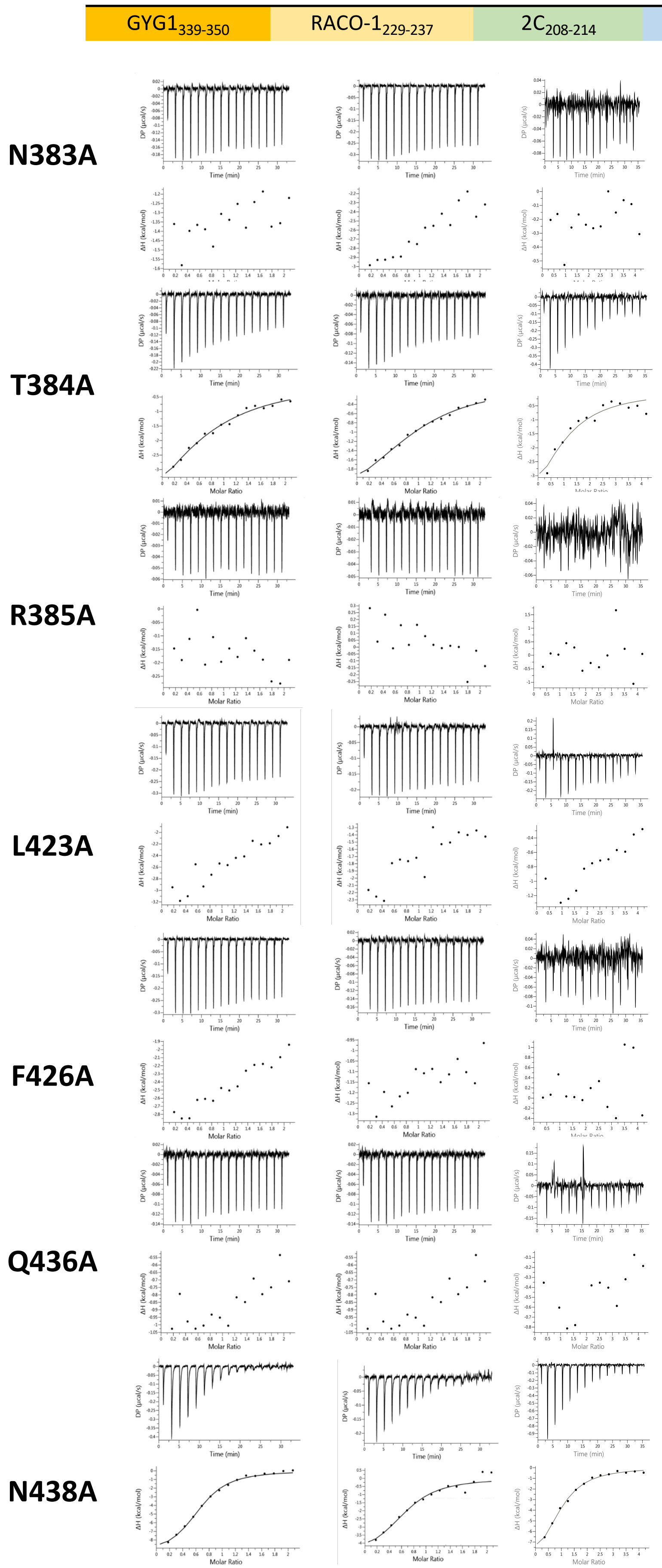

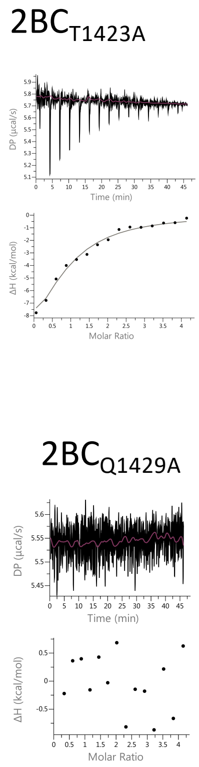

| Trim7 | GYG1$_{339-350}$ | RACO-1$_{229-237}$ | 2BC$_{1423-1429}$ | NS6$_{1171-1177}$ |
|---|---|---|---|---|
| WT | 8 ± 0.6 µM | 7.9 ± 1.0 µM | 11.2 ± 4.2 µM | 8.7 ± 3.2 µM |
| N383A | No Binding | No Binding | No binding | NT |
| T384A | 22.6 ± 6.6 µM | 12.3 ± 2.0 µM | 21.2 ± 5.84 µM | NT |
| R385A | No Binding | No Binding | No binding | No binding |
| L423A | No Binding* | No Binding* | No binding* | NT |
| F426A | No Binding | No Binding | No binding | NT |
| Q436A | No Binding | No Binding | No binding | NT |
| N438A | 2.3 ± 0.3 µM | 3.9 µM ± 1.8 µM | 7.82 ± 1.18 µM | NT |

**Figure S4.** GYG isoforms

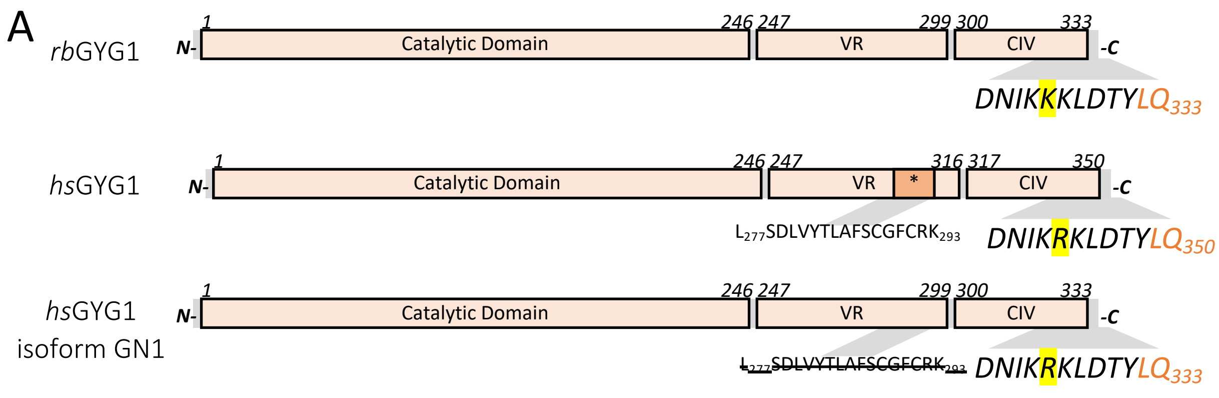

**Figure S5.** Stable cell line validation.

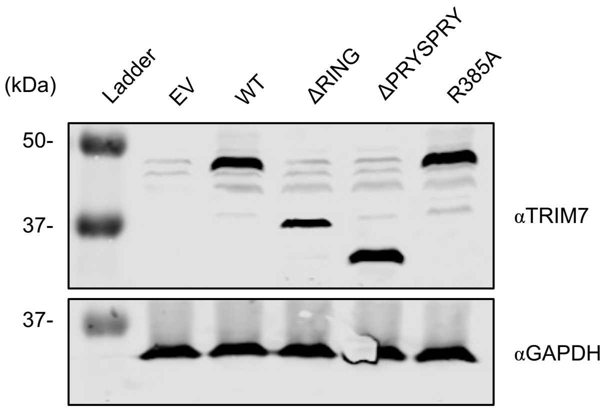

**Figure S6.** SARS-CoV-2 infection is not restricted by TRIM7

A

| Protein | C-term |
|---|---|
| NSP4 | 'LQ' |
| NSP5 | 'FQ' |
| NSP6 | 'VQ' |
| NSP7 | 'LQ' |
| NSP8 | 'LQ' |
| NSP9 | 'LQ' |
| NSP10 | 'LQ' |
| NSP11 | |
| NSP12 | 'LQ' |
| NSP13 | 'LQ' |
| NSP14 | 'LQ' |
| NSP15 | 'LQ' |

B

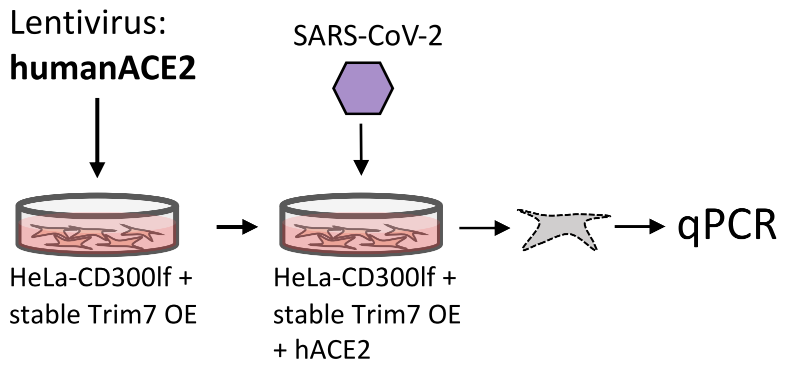

C

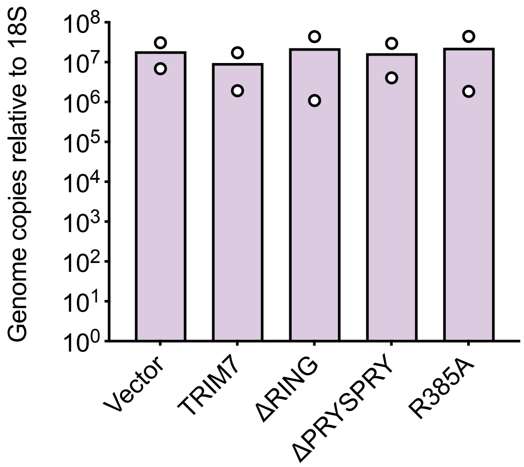

### **Figure S7.** Phylogenetic trees for GYG1 and TRIM7

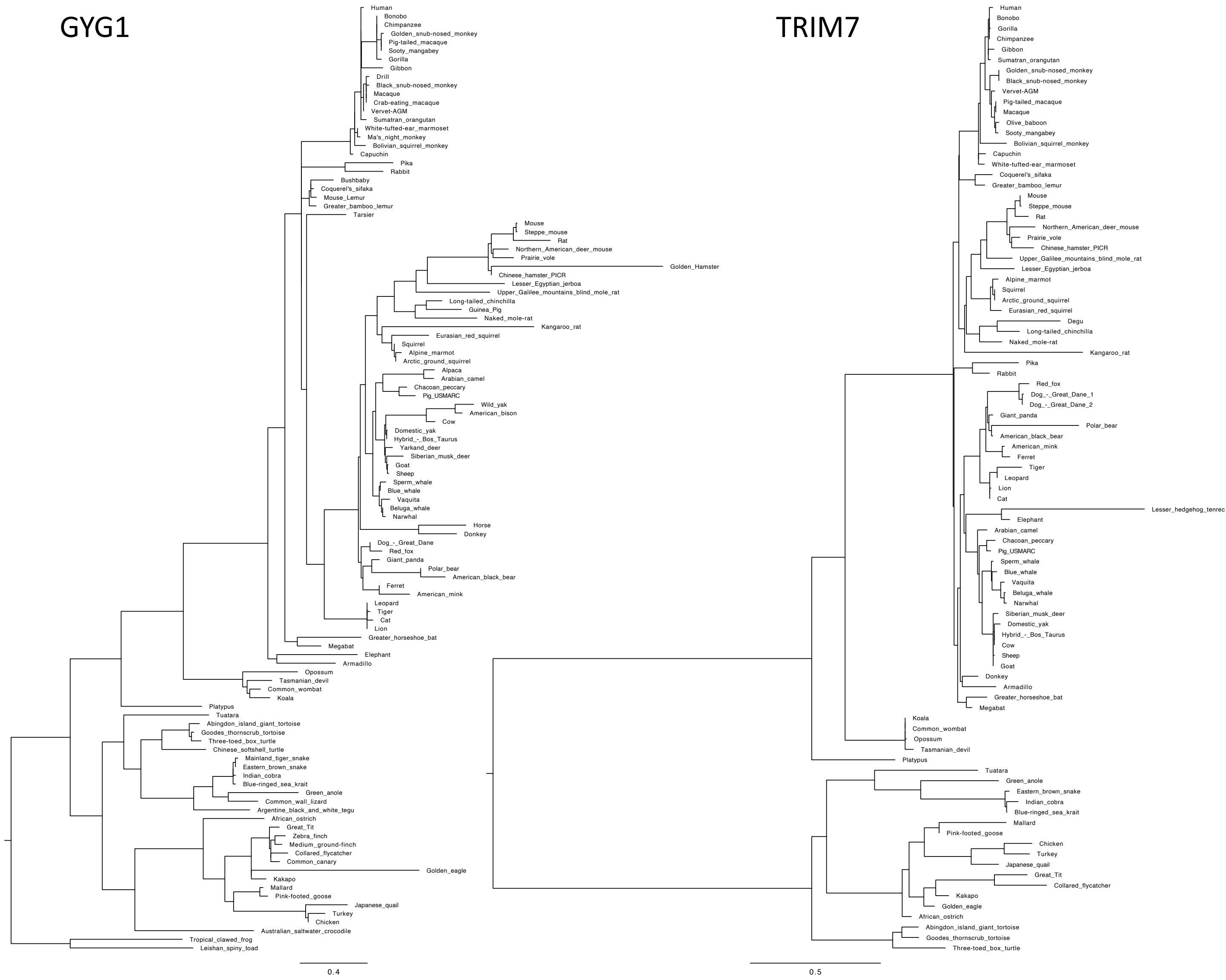

#### Table S1 Data collection and refinement statistics

| | 7OW2 (TRIM7: GYG1$_{322-333}$) | 7OVX (TRIM7: RACO-1$_{229-235}$) | 8A5L (TRIM7:2C$_{208-214}$) | 8A5M (TRIM7:MNV1-NS6$_{1171-1177}$) | 8A8X (TRIM7:MNV1-NS3$_{699-705}$) |
|---|---|---|---|---|---|
| **Data collection** | | | | | |
| Space group | P212121 | P61 | P65 | P1211 | P1211 |
| Cell dimensions | | | | | |
| *a*, *b*, *c* (Å) | 37.63, 54.23, 81.40 | 108.12, 108.12, 137.14 | 79.89, 79.89, 53.2 | 40.92, 112.97, 53.04 | 51.25, 53.67, 73.60 |
| α, β, γ (°) | 90.0, 90.0, 90.0 | 90.0, 90.0, 120.0 | 90.0, 90.0, 120.0 | 90.0, 102.93, 90 | 90, 105.5, 90 |
| Resolution (Å) | 19.3-1.7 (1.78-1.70) | 93.64-2.2 (2.29-2.20) | 29-1.62 (1.62-1.66) | 112.97-2.92 (3.10-2.92) | 70.91-2.37 (2.37-2.43) |
| $R_{meas}$ | 9.6 (30.1) | 10.3 (48.1) | 11.4 (92.5) | 9.4 (N/A) | 17.7 (96.1) |
| $CC_{1/2}$ (%) | 99.4 (91.2) | 99.7 (96.3) | 99.7 (59.9) | 96.4(33.8) | 100 (60) |
| $I / \sigma I$ | 8.9 (2.8) | 12.6 (3.1) | 11.1 (1.6) | 24.4(1.4) | 6.8 (1.2) |
| Completeness (%) | 99.0 (98.8) | 99.2 (98.5) | 99.7 (95.6) | 99.9 (98.8) | 97.4 (74.4) |
| Redundancy | 3.3 (3.1) | 6.4 (5.6) | 4.9 (3.7) | 3.5 (3.4) | 3.3 (2.4) |
| Resolution (Å) | 1.7 | 2.2 | 1.62 | 2.92 | 2.37 |
| No. reflections | 17598 | 47982 | 24565 | 10246 | 15418 |
| $R_{work}$ / $R_{free}$ | 0.18/0.22 | 0.20/0.24 | 0.15/0.18 | 0.19/0.29 | 0.19/0.26 |
| No. atoms | 1657 | 5750 | 1619 | 2857 | 2875 |
| Protein | 1506 | 5614 | 1443 | 2815 | 2819 |
| Ligand/ion | 0 | 2 | 10 | 15 | 0 |
| Water | 151 | 135 | 152 | 24 | 56 |
| *B*-factors | | | | | |
| Protein | 21.81 | 41.62 | 17.99 | 57.69 | 30.68 |
| Ligand/ion | 0 | 46.39 | 33.52 | 92 | N/A |
| Water | 33.54 | 36.74 | 30.03 | 33.6 | 26.02 |
| R.m.s. deviations | | | | | |
| Bond lengths (Å) | 0.01 | 0.01 | 0.01 | 0.01 | 0.01 |
| Bond angles (°) | 1.60 | 1.40 | 1.7 | 1.9 | 1.7 |

*Values in parentheses are for highest-resolution shell.

**Table S2 – ITC data.** Concentration for proteins and peptides is given as monomer. * indicates where N was fixed in analysis. **Concentration of GYG1 (and MBP-T7-CC-PS) dimer was used in the analysis. NB = no binding.

| TRIM7 | CONC. [UM] | LIGAND | CONC. [UM] | SEQUENCE | N | $K_D$ | ΔH (CAL/MOL) | TEMP[°C] |
|---|---|---|---|---|---|---|---|---|
| **HTR7-PS WT** | 500 | rbGYG1 | 100 | see methods | 1.07±0.014 | 2.2 µM±0.17 | -7271±130 | 15 |
| **MTR7-PS WT** | 500 | rbGYG1 | 100 | see methods | 0.93±0.005 | 1.5 µM±0.27 | -1113±79 | 15 |
| **MTR7-PS WT** | 500 | mGYG1 | 100 | see methods | 0.96±0.008 | 1.04 µM ±0.15 | -1192±132 | 15 |
| **HTR7-PS WT** | 20 | hGYG1-GN1 | 400 | see methods | 0.42±0.03 ** | 2.16 µM±0.06 | -27000±3300 | 25 |
| **HTR7-PS WT** | 500 | rbGYG1$_{VR-CIV}$ | 100 | see methods | 1.05±0.011 | 2.6 µM ±0.33 | -6433±95 | 15 |
| **HTR7-PS WT** | 500 | rbGYG1$_{CIV}$ | 100 | see methods | 0.70±0.034 | 4.8µM ±4.0 | -9191±615 | 15 |
| **HTR7-PS WT** | 500 | rbGYG$_{322-333}$ | 50 | DNIKKKLDTYLQ | 0.70±0.03 | 21 ± 1.8 µM | -9191±615 | 15 |
| **HTR7-PS WT** | 25 | hGYG$_{338-350}$ | 250 | DNIKRKLDTYLQ | 0.97±0.02 | 8 ± 0.64 µM | -11200±383 | 20 |
| **HTR7-PS WT** | 25 | RACO-1$_{227-235}$ | 250 | NLGLSMLQ | 0.61±0.04 | 7.9 ± 1.06 µM | -6360±561 | 20 |
| **HTR7-PS WT** | 20 | CVB-2C$_{208-214}$ | 400 | TIEALFQ | 0.90±0.2 | 11.2 ± 4.2 µM | -12600±3580 | 25 |
| **HTR7-PS WT** | 20 | MNV1-NS6$_{1172-1178}$ | 400 | LEALEFQ | 1.13±1.62 | 8.7 ± 3.2 µM | -11700±2250 | 25 |
| **HTR7-PS WT** | 20 | MNV1-NS3$_{700-706}$ | 400 | HDDFGLQ | 1.18±0.19 | 14.3 ± 4.3 µM | -10900±2280 | 25 |
| **HTR7-PS WT** | 20 | polyA | 400 | AAAAAAA | NB | NB | NB | 25 |
| **HTR7-PS WT** | 20 | polyA-LQ | 400 | AAAAALQ | 1* | 8.92 ± 3.03 µM | -4870±513 | 25 |
| **HTR7-PS WT** | 20 | LQ | 2000 | LQ | 1* | 201 ± 30 µM | -5720±507 | 25 |
| **HTR7-PS WT** | 20 | Ac-LQ | 800 | Ac-LQ | 1* | 144 ± 26 µM | -6600±760 | 25 |
| **HTR7-PS WT** | 20 | Ac-LLQ | 800 | Ac-LLQ | 1* | 38.7 ± 5.16 µM | -7720±484 | 25 |
| **HTR7-PS WT** | 20 | TIEALFA | 400 | TIEALFA | NB | NB | NB | 25 |
| **HTR7-PS WT** | 20 | AIEALFQ | 400 | AIEALFQ | 0.80±0.19 | 17.0 ± 4.3 µM | -15500±4410 | 25 |
| **HTR7-PS N383A** | 25 | hGYG$_{338-350}$ | 250 | DNIKRKLDTYLQ | NB | NB | NB | 20 |
| **HTR7-PS T384A** | 25 | hGYG$_{338-350}$ | 250 | DNIKRKLDTYLQ | 0.71±0.18 | 22.6 ± 6.6 µM | -7280±2410 | 20 |
| **HTR7-PS R385A** | 25 | hGYG$_{338-350}$ | 250 | DNIKRKLDTYLQ | NB | NB | NB | 20 |
| **HTR7-PS L423A** | 25 | hGYG$_{338-350}$ | 250 | DNIKRKLDTYLQ | NB | NB | NB | 20 |
| **HTR7-PS F426A** | 25 | hGYG$_{338-350}$ | 250 | DNIKRKLDTYLQ | NB | NB | NB | 20 |
| **HTR7-PS Q436A** | 25 | hGYG$_{338-350}$ | 250 | DNIKRKLDTYLQ | NB | NB | NB | 20 |
| **HTR7-PS N438A** | 25 | hGYG$_{338-350}$ | 250 | DNIKRKLDTYLQ | 0.64±0.02 | 2.32 ± 0.26 µM | -9830±318 | 20 |
| **HTR7-PS N383A** | 25 | RACO-1$_{227-235}$ | 250 | NLGLSMLQ | NB | NB | NB | 20 |
| **HTR7-PS T384A** | 25 | RACO-1$_{227-235}$ | 250 | NLGLSMLQ | 0.99±0.05 | 12.3 ± 2.00 µM | -2870±253 | 20 |
| **HTR7-PS R385A** | 25 | RACO-1$_{227-235}$ | 250 | NLGLSMLQ | NB | NB | NB | 20 |
| **HTR7-PS L423A** | 25 | RACO-1$_{227-235}$ | 250 | NLGLSMLQ | NB | NB | NB | 20 |
| **HTR7-PS F426A** | 25 | RACO-1$_{227-235}$ | 250 | NLGLSMLQ | NB | NB | NB | 20 |
| **HTR7-PS Q436A** | 25 | RACO-1$_{227-235}$ | 250 | NLGLSMLQ | NB | NB | NB | 20 |
| **HTR7-PS N438A** | 25 | RACO-1$_{227-235}$ | 250 | NLGLSMLQ | 0.67±0.08 | 3.93 ± 1.75 µM | -4930±852 | 20 |
| **HTR7-PS N383A** | 20 | CVB-2C$_{208-214}$ | 400 | TIEALFQ | NB | NB | NB | 25 |
| **HTR7-PS T384A** | 20 | CVB-2C$_{208-214}$ | 400 | TIEALFQ | 1 * | 21.2 ± 5.84 µM | -6120±704 | 25 |
| **HTR7-PS R385A** | 20 | CVB-2C$_{208-214}$ | 400 | TIEALFQ | NB | NB | NB | 25 |
| **HTR7-PS L423A** | 20 | CVB-2C$_{208-214}$ | 400 | TIEALFQ | NB | NB | NB | 25 |
| **HTR7-PS F426A** | 20 | CVB-2C$_{208-214}$ | 400 | TIEALFQ | NB | NB | NB | 25 |
| **HTR7-PS Q436A** | 20 | CVB-2C$_{208-214}$ | 400 | TIEALFQ | NB | NB | NB | 25 |
| **HTR7-PS N438A** | 20 | CVB-2C$_{208-214}$ | 400 | TIEALFQ | 0.92±0.06 | 7.82 ± 1.18 µM | -10400±910 | 25 |
| **HTR7-PS R385A** | 20 | MNV1-NS6$_{1172-1178}$ | 400 | LEALEFQ | NB | NB | NB | 25 |
| **HTR7-PS R385A** | 20 | MNV1-NS3$_{700-706}$ | 400 | HDDFGLQ | NB | NB | NB | 25 |
| **HTR7-CC-PS WT** | 12 | hGYG$_{338-350}$ | 300 | DNIKRKLDTYLQ | 1.7 ± 0.8 | 15 ± 6 µM | -9680±5770 | 25 |
| **HTR7-CC-PS WT** | 12 | hGYG1-GN1 | 150 | see methods | 0.8 ± 0.02 | 0.16 ± 0.05 µM | -23800±991 | 25 |
